# An assessment of bioinformatics tools for the detection of human endogenous retroviral insertions in short-read genome sequencing data

**DOI:** 10.1101/2022.02.18.481042

**Authors:** Harry Bowles, Renata Kabiljo, Ashley Jones, Ahmad Al Khleifat, John P Quinn, Richard JB Dobson, Chad M Swanson, Ammar Al-Chalabi, Alfredo Iacoangeli

## Abstract

There is a growing interest in the study of human endogenous retroviruses (HERVs) given the substantial body of evidence that implicates them in many human diseases. Although their genomic characterization presents numerous technical challenges, next-generation sequencing (NGS) has shown potential to detect HERV insertions and their polymorphisms in humans, and a number of computational tools to detect them in short-read NGS data exist. In order to design optimal analysis pipelines, an independent evaluation of the currently available tools is required. We evaluated the performance of a set of such tools using a variety of experimental designs and types of NGS datasets. These included 50 human short read whole-genome sequencing samples, matching long and short read NGS data, and simulated short-read NGS data. Our results highlight the performance variability of the tools across the datasets and suggest that different tools might be suitable for different study designs. Using multiple tools and a consensus approach is advisable if computationally feasible and wet-lab validation via PCR is advisable where biological samples are available.

## Introduction

Endogenous retroviruses (ERVs) integrated into the genome of vertebrates as a result of ancient exogenous infections. They invaded the germ cell lines of all vertebrates including humans, becoming an integral part of the germline transmission and therefore replicate in a Mendelian fashion (1). Human endogenous retroviruses (HERVs) comprise ~8% of the genome, whereas protein coding genes comprise only 1-2% (2). Although they make up a striking portion of the human genome, most of them are inactive as a consequence of the accumulation of mutations and DNA methylation (3). The HML-2 HERV-K subgroup includes some of the most recent HERV integrations, which are found as full length (or near full length) insertions in over 80 different loci (4). Though there are several full-length copies of HERV-K in the genome, none are likely to produce an infectious virus (5). HERV-K sequences can be full length proviruses, solo long terminal repeats or 2-LTR sequences and are polymorphic in the human population (6). A full length HERV-K provirus consists of long terminal repeats (LTRs) flanking the viral genes (*gag, pro, pol* and *env*). In the majority of elements defined as HERV loci in the human genome, only the LTRs are present which contain the promoter and enhancer regions (7) (figure 1).

**Figure 1).**
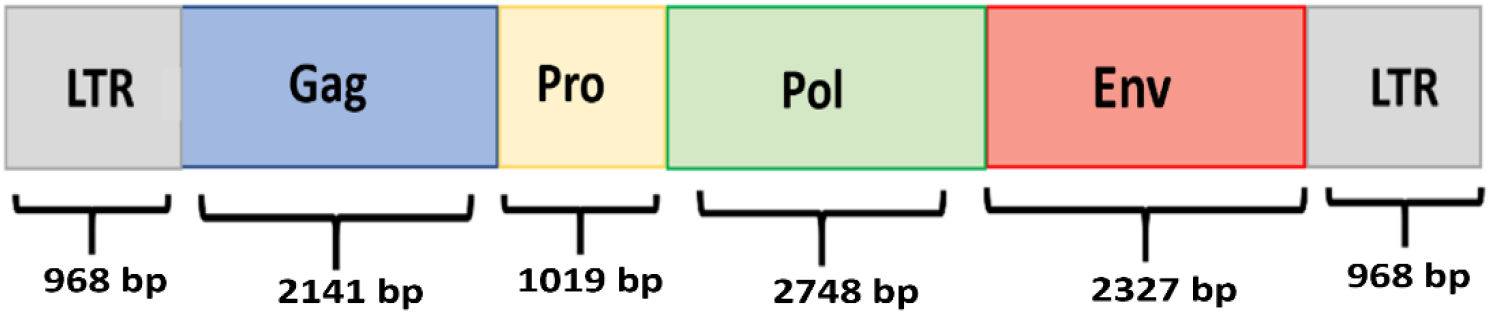
This schematic shows the general structure of a full length HERV-K with approximate base pair lengths for each section. The LTR length is the LTR5_Hs length given in the DFAM database (accession = DF0000558), the internal gene lengths comes from HERVK11 internal regions, again from DFAM (accession = DF0000189) (8). The *gag* gene encodes a polyprotein that is cleaved to produce the structural proteins that make up a viral particle. The *pro* gene encodes the viral protease while *pol* codes for the viral enzymes required for reverse transcription and integrase. The *env* gene codes for the viral envelope protein that mediates viral entry into a target cell. The gene products require post translational cleavage, for example, *pro* cleaves the *gag* polyprotein into ‘matrix’, ‘capsid’ and ‘nucleocapsid’ subunits. The majority of HERV-K loci only contain the LTR regions which contain the enhancer and promoter sequences that regulate transcription.

HERVs, and related transposable elements, have been linked to a wide range of diseases including cancer and neurodegeneration via multiple mechanisms. For example, their insertion into the human genome may alter gene expression or disrupt reading frames (9); they were reported to be upregulated in biological samples from people affected by neurodegenerative diseases and cancer (10,11,12). Furthermore, their expression may be toxic for certain cell types such as motor neurons (13). However, characterising the HERV genomic landscape is challenging. This is due to several factors including some characteristics of their sequence which make them hard to map correctly in the human genome. For example, HERVs are thousands of bases long, they are polymorphic in humans and present a high degree of sequence similarity. As a consequence, despite the substantial body of evidence supporting their potential key role in many diseases, they remain substantially under-investigated and the nature of their involvement with human disease is not clear.

Recent advances in next-generation sequencing (NGS) have made sequencing DNA molecules a common practice in genetic research, allowing for the investigation of a wide range of variants that spans from single nucleotide variants to large structural variants (14). This technology has also been used to study HERVs in humans (15). Currently, a number of bioinformatic tools for the identification of HERV loci in NGS data exist, mostly based on the exploitation of split and discordant reads to reveal the presence of potential HERV insertions (Figure 2). Given the growing interest in this class of elements, an independent evaluation of the current tools for HERV detection in greatly needed for the design of optimal analyses pipelines and to promote the discussion necessary for the scientific community to establish best practice protocols. On this basis, we designed a set of experiments to benchmark widely used computational tools and protocols for the detection of HERVs in short-read NGS data (short reads = 50 - 200bp). Considering their proposed role in human disease, we focused our experiment on the identification of HERV-K insertions that are not present in the human reference genome (non-reference HERV-Ks). We tested six widely used tools on three short-read sequencing datasets: a large short-read whole-genome sequencing (SR-WGS) dataset (50 samples), a simulated SR-WGS dataset, and six SR-WGS samples for which matching long-read sequencing data was available.

**Figure 2).**
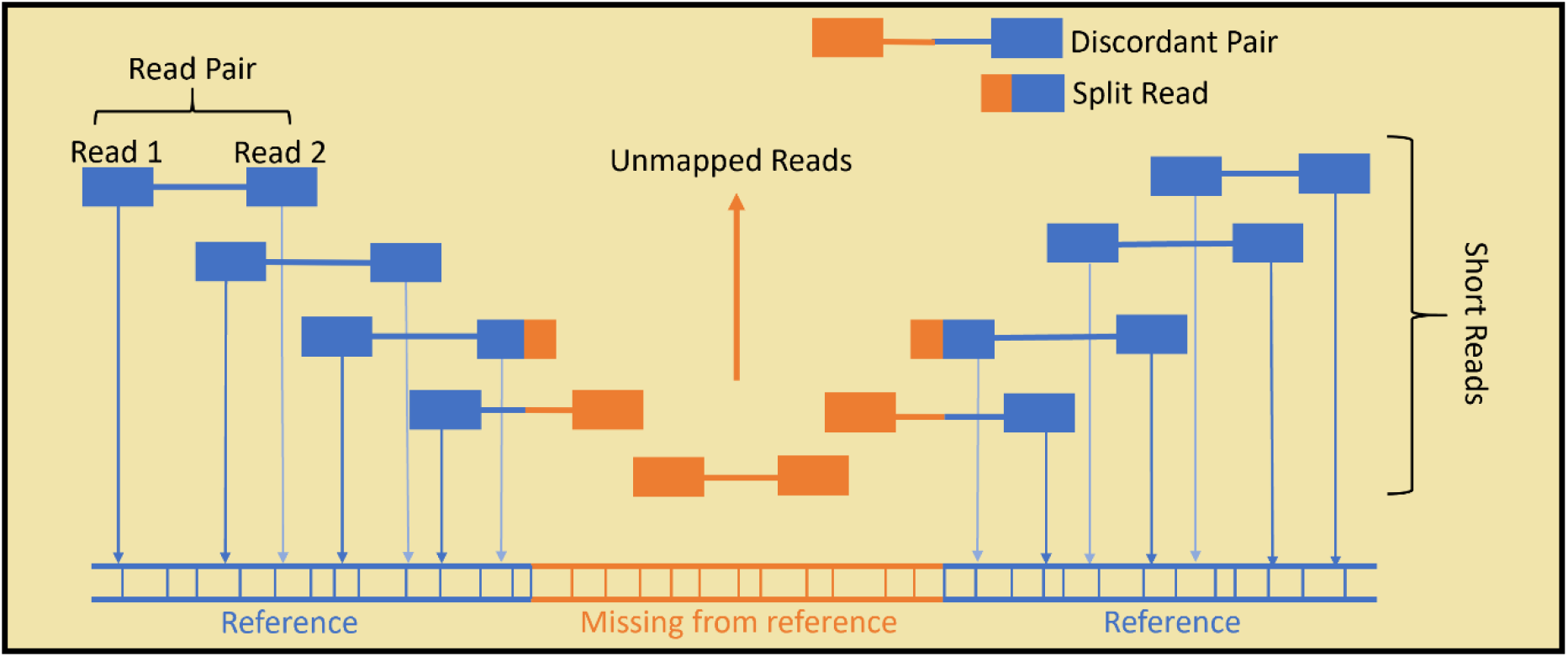
In standard Illumina paired end SR-WGS, the DNA sample is first sheared into small fragments. Then, both ends of the fragment are sequenced, giving paired reads. Read mapping is achieved by aligning the reads to a reference genome. HERVs that are present in the sample, but not the reference, cannot be aligned fully and remain largely unmapped – which leads them to often be ignored in genome analysis. Specialised bioinformatics tools use unmapped, split and discordant reads to predict non-reference HERVs.

## Methods

### Overview of the tested tools

#### MELT

MELT scans WGS data for clusters of discordant read pairs and split reads. Split and discordant reads can be mapped to an insertional element reference sequence provided by the user, to allow the detection of specific insertion types. It can also genotype reference mobile elements (16). According to its documentation, MELT was not tested for HERV detection, and the authors predicted that it may perform poorly on LTR elements compared to non-LTR transposable elements (such as Alus). Nevertheless, MELT has been used to detect HERVs (19,20,22).

#### Mobster

This tool also uses discordant reads alongside split reads and an insertional reference sequence to predict specific insertion sites. When Mobster was released, the authors reported that it was not able to identify HERV insertions. However, they tested it using two WGS paired-end samples and since its first publication in 2014, Mobster has been extensively upgraded and gained a considerable popularity. We included it in our experiment, tested its updated version on a larger sample and used a different HERV-K template sequence to that used in the authors’ benchmarking work (17).

#### Retroseq

Retroseq uses discordant read pairs to identify putative insertions sites and filters for read pairs which align to a reference of interest (18).

#### Steak

Steak annotates both reference and non-reference mobile elements. Unlike the other tools, it first identifies reads that partially map to the target HERV reference sequence. These reads are assumed to map to the edge of the insertion. These mapped fragments are removed, and a library of host reference flank reads and their mates, is created. These reads are mapped onto the human reference genome to identify the presence of both reference and non-reference HERV loci (19).

#### ERVcaller

The tool extracts incorrectly mapped read pairs and split reads to identify likely insertions sites, that are then aligned to a reference sequence to allow for the detection of specific insertion types (20).

#### Retroseq+

This pipeline is our in-house implementation of a protocol described by Wildschutte et al. It uses Retroseq as a base for predicting HERV-K insertion loci. It then refines the results through insertion junction reconstruction and secondary scanning of the junction for HERV-K sequence by RepeatMasker. Because this protocol is not available as a automatic bioinformatics pipeline, we have implemented it ourselves, following the authors’ description (21).

##### Benchmarking experiments

In order to assess the performance of these tools, we set up four experiments: i) using the tools on 50 SR-WGS samples, we measured the agreement between tools; ii) using the HERV calls from the 50 SR-WGS samples, we attempted to validate the tools by quantifying the proportion of predicted HERV-K insertions that were previously reported in literature; iii) we assessed the specificity of each tool by using long-read data for the validation of HERV-K calls from matching short-read data; iv) finally, we estimated the performance of the tools using simulated NGS data.

##### Overlap analysis and comparison with previously reported HERVs

Each tool was applied to WGS data of 50 amyotrophic lateral sclerosis (ALS) patients from the British Project Mine dataset (23,24). This WGS was generated by sequencing DNA samples from blood, using the Illumina Hiseq 2000 platform. The resulting WGS samples had read length equal to 100bp with an average coverage depth of 40X (paired end reads). We aligned them to the hg19 reference genome using Burrows-Wheeler alignment, BWA-MEM (25). The predicted insertion sites across all genomes were compared to a list of 40 well characterised polymorphic HERV-K insertions previously described in the literature [S1] (26). They were also compared to all reference HERV loci (both HERV-Ks and wider HERV/LTR subgroups) obtained through the UCSC table browser using the RepeatMasker (RMSK) track for the hg19 genome build. The following identifiers were used to retrieve HERV-K reference loci: LTR5_Hs, LTR5A, LTR5B, HERV-K and HERV-K-int [S2]; while the set of all reference HERVs was obtained by taking the entire UCSC RMSK hg19 track and extracting those which had the identifiers ‘ERV’ or ‘LTR’ [S3]. HML-2 type HERV-Ks, which are targeted in this analysis can be subclassified based on their LTR sequence. LTR5B is the phylogenetically oldest HML-2 LTR, LTR5_Hs is younger and human specific.

An overlap was defined as a predicted insertion being within 500 base pairs of a known ERV locus. The number of reference HERV-Ks and HERVs per million bases across the human chromosomes is shown in Figure 3.

**Figure 3).**
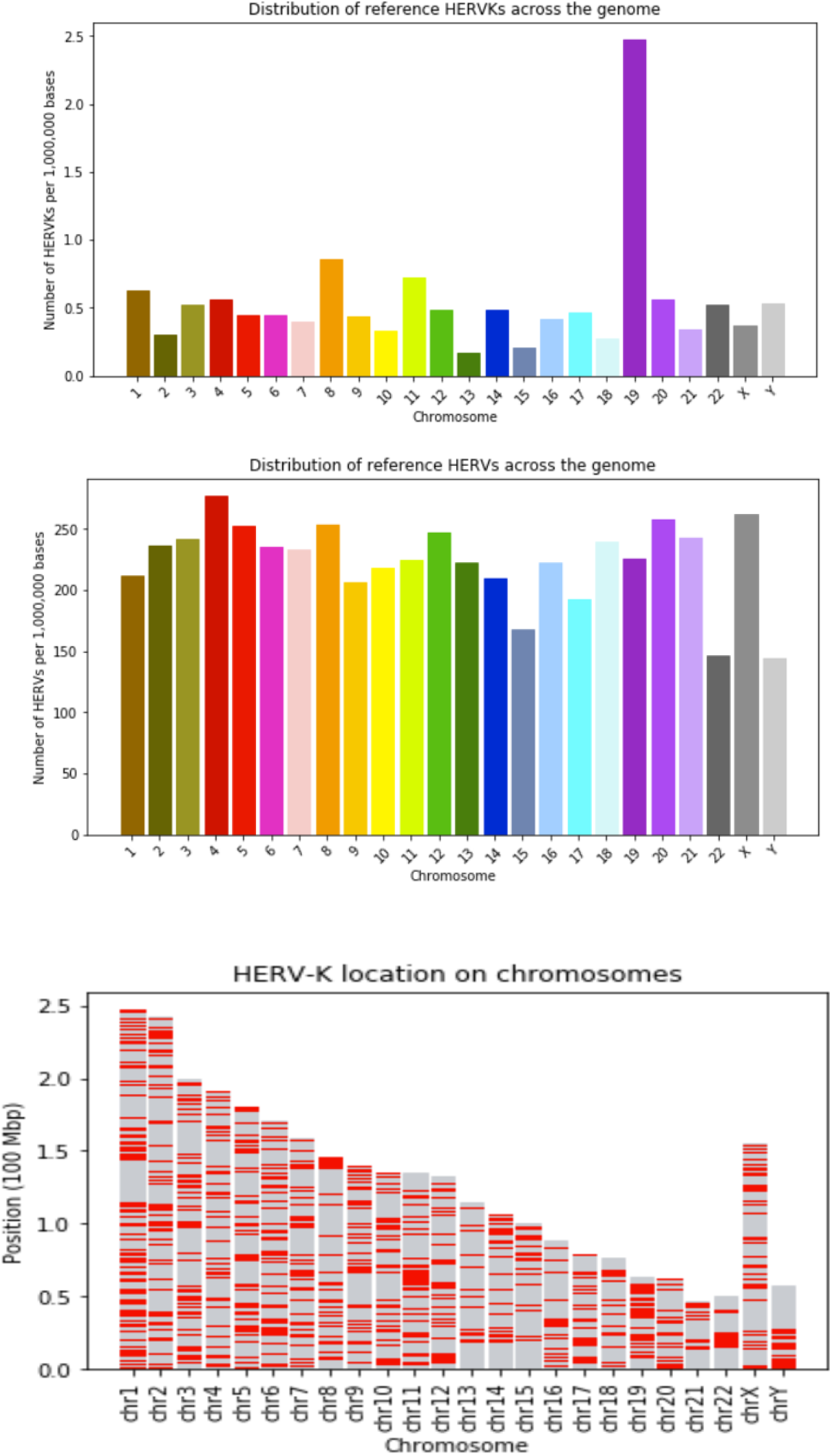
Upper panel - Number of HERV-Ks per million bases in each human chromosome (as given by the UCSC RMSK table). The high HERV density in chr19 has been previously reported (27) and other transposable elements are also enriched on this chromosome (28). Middle panel - Number of HERVs and LTR sequences per million bases in each human chromosome – Data is from the UCSC RMSK table. Lower panel – this panel shows the distribution of HERV-K on each chromosome, with a red line indicating the presence of the HERV-K LTR. These results are obtained from the LTR5/HERV-K UCSC RMSK table.

The results of each tool were also compared to one another, giving for each tool, the proportion of its results that were also predicted by each of the other tools. This allowed us to quantify the agreement across tools. For Steak and Melt, reference results were filtered out from the total results using the UCSC RMSK table of hg19 reference HERV-K loci [S2].

##### Long-read sequencing data

We used a set of six samples from Wang et al, for which both short and long-read genome sequencing data were available (29). Briefly, the short-read data (derived from blood) were sequenced using Illumina Hiseq 2500 giving mean 105bp paired end reads with coverage depth ranging between 15.6X and 18.8X. The long reads were sequenced using PacificBio Seqel system version 2. For these samples the read lengths were between 10KB and 18KB and the mean coverage depth of samples ranged between 28.5X and 69X.

Long-read sequencing (read length > 10,000 base-pairs) can capture a large overhang, if not the whole, of the HERV-K allowing for their accurate identification (30,31,32). We applied each tool to the short-read WGS data to predict LTR5_Hs HERV-K insertions and used the long-read data for validation as follows. For each predicted insertion, we extracted long reads mapped at the corresponding locus from the matching long-read WGS sample. These long reads were then assembled into contigs using wtdbg2 (33) and RepeatMasker (34) was applied to the contigs to confirm the presence of a HERV-K LTR sequence at this site. For Melt and Steak, reference HERV-Ks were filtered out.

##### Simulated SR-WGS analysis

For the simulation test, short-read paired-end Illumina WGS data were simulated from the hg19 reference sequence using DWGSIM (parameters in Table 1) (35).

**Table 1).**
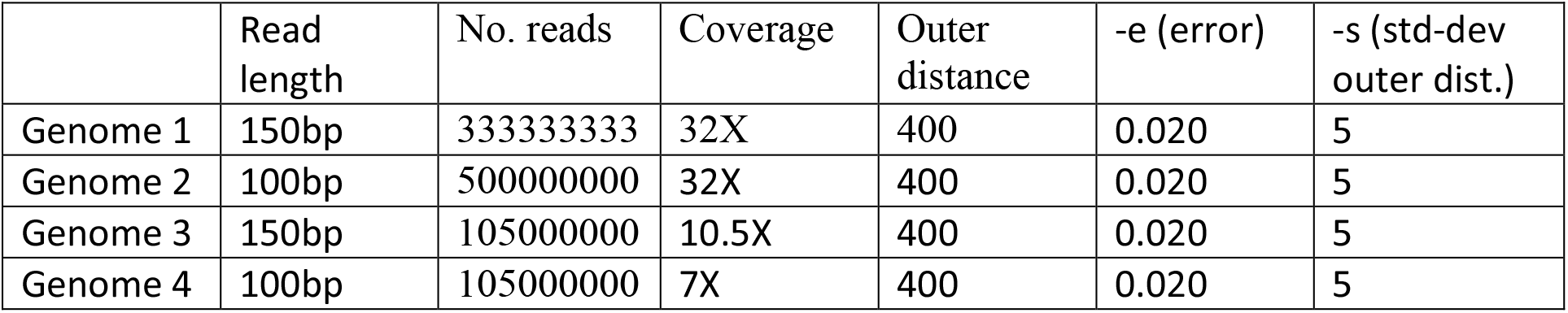
Custom parameters used to generate the simulated data. All parameters not included in this table were kept as default.

Hg19 is reported to have 66 HERV-K (HML-6) proviral loci (2) from which we selected 19. To simulate these 19 HERV-K sites as novel insertions, after generating the WGS data, we removed the 19 sequences from hg19 using Bedtools masking followed by deletion of the mask (36). The generated FASTQ files were then aligned to the edited hg19 using BWA-MEM. Thus, the simulated data contained 19 known HERV-K insertions that were not present in our edited reference genome, and these 19 insertions were the only non-reference elements in the simulated samples. Each tool was then applied to the simulated WGS, and the 19 sites were used as our set of true positive insertions. 15 insertions were type LTR3A and 4 were type LTR3B. The LTR3A sequence was used as target reference template. This allows us to see how well each tool can distinguish specific insertion types as well as assess general sensitivity.

We defined sensitivity as the proportion of the known, non-reference insertions which were successfully detected: True positive/(True positive + False negative). We defined precision as the proportion of positive results that are true: True positive/(True positive + False positive).

##### Computational efficiency report

We tested the memory usage and time taken for each tool to run on a single WGS sample from the Project MinE dataset. Slurm was the scheduling system on the Linux HPC platform used for this project. Slurm has its own command (sacct) for timing scripts and assessing memory usage. Each tool was applied to a single WGS sample from Project MinE and sacct was used to report the tools memory and cpu usage. To determine the sizes of intermediate files produced by each tool, the ‘du’ command was run on a loop, executing every second, over the directory in which each tool was running. The difference between the starting directory size and largest directory size reported by ‘du’ is reported.

## Results

### Analysis of the 50 short-read WGS samples

Each tool was applied to 50 SR-WGS samples and the results were merged. Table 2 shows the proportion of predicted HERV-K loci that map to a known HERV loci. Total number of predictions and overlap with previously reported loci greatly varied across tools (Table 2 and Figure 4). Results varied greatly across tools. Notably, Steak gave the highest number of predictions and 84% of these results matched to previously documented HERV locations. 97.6% of the results given by Retroseq+ matched to a documented HERV loci though this tool gave the smallest number of predictions. Mobster was not able to detect any HERV-Ks in this sample.

**Figure 4).**
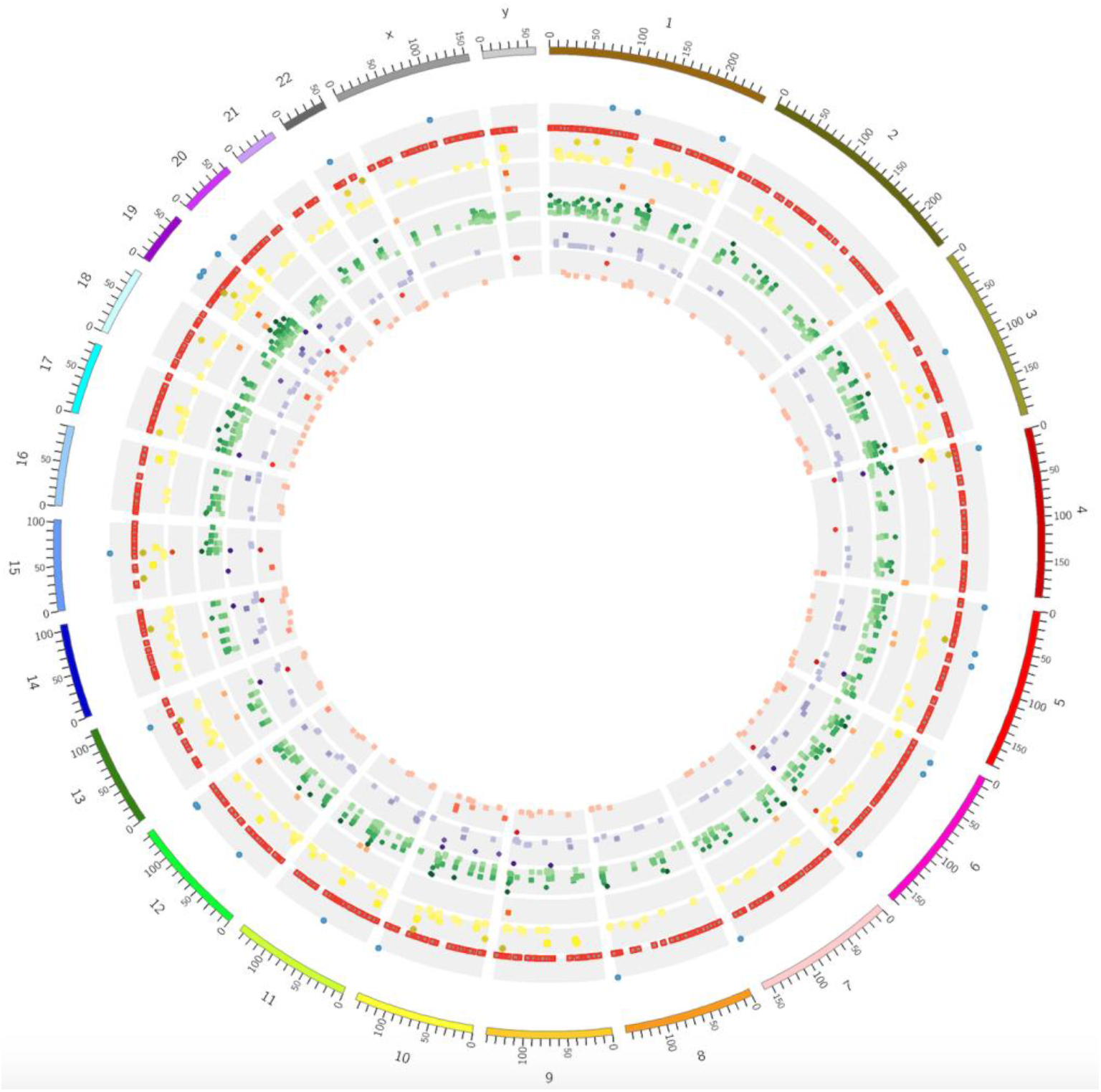
Overview of novel insertions predicted by the tools on the 50 genomes in a circular chromosomal plot. The order of concentric circles from the outside of the plot: circle 1 (blue dots) – known non reference insertions; circle 2 (red dots) – known reference insertions; circle 3 (yellow): Retroseq predictions; circle 4 (orange): Retroseq plus predictions; circle 5 (green): Steak predictions; circle 5 (purple) ERVcaller predictions, circle 6 (red): Melt predictions. The intensity of the colour and the height of each dot in its band is proportional to the number of subjects in whom the insertion is predicted with darker colours and higher position corresponding to a larger number.

**Table 2).**
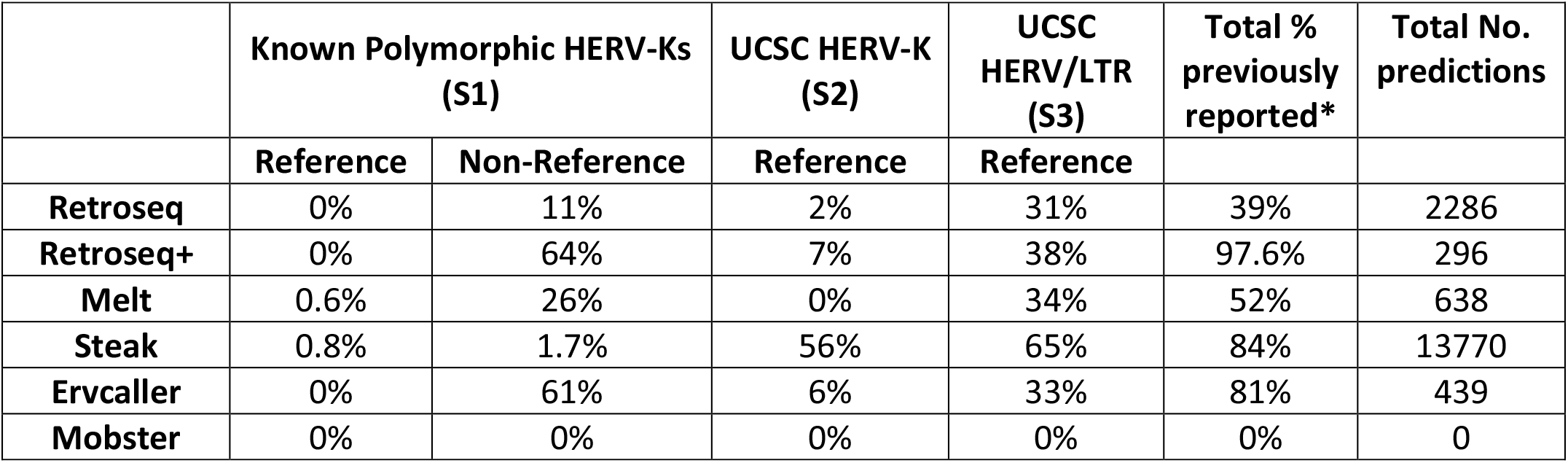
The ‘Known Polymorphic’ column shows the proportion of predicted HERV-K insertion loci that matched to HERV-Ks from the literature reported to be polymorphic [S1]. UCSC HERV-K and UCSC HERV columns show the proportion which matched to hg19 reference HERV-Ks and HERVs given by the UCSC table browser [S2-S3]. Total percent previously reported shows the proportion of predictions that are present in the polymorphic set or in the UCSC sets. Total No. predictions is the total number of predictions given across all 50 genomes. *There is an overlap between the Non-reference polymorphic HERV-Ks and the UCSC HERV/LTR set, this explains why the ‘Total % previously reported’ column is less than a sum of these two columns for most tools.

The agreement between tools (Figure 5) greatly varied, ranging between 2.8% (proportion of Steak calls that were also called by Melt) and 63% (proportion of Retroseq+ calls that were also called by Steak).

**Figure 5).**
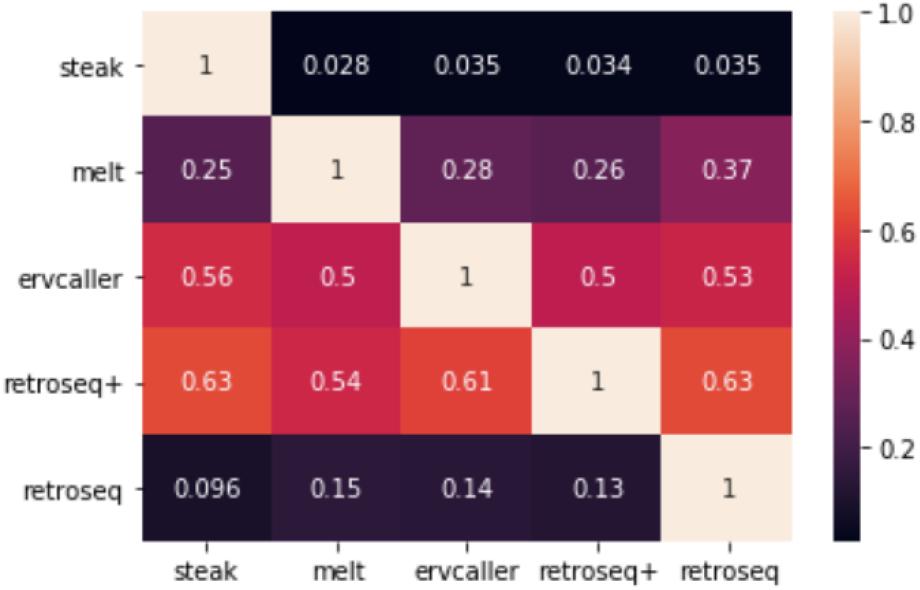
Heatmap reporting the proportion of insertions found by tools on the rows that were also found by the tools on the columns. E.g. the proportion in [Steak, Ervcaller] represents the proportion of Steak calls that were also called by Ervcaller. Therefore, please note that [Steak, Ervcaller] is not equal to [Ervcaller, Steak]. Reference HERV-Ks have been filtered out.

### Simulated data results

Each tool was applied to a set of four simulated WGS samples, with known HERV insertions of different types (LTR3A and LTR3B). Each tool was given LTR3A as target element. All tools performed worse on samples with lower read depth. For example, Retroseq detected 11/15 LTR3A insertions in a 32X sample but only 7/15 LTR3A insertions in a 10.5X and 7X sample. However, the degree to which read depth affects each tool varied. ERVcaller performed equally well on 32X and 10.5X sample finding 11 of the 15 LTR3A insertions and, in the 7X sample, it still identified 10 insertions. The only tool to mistakenly detect an LTR3B insertion was Retroseq. Sensitivity and precision greatly varied across tools. E.g. Steak had the highest average precision (0.9) while Retroseq+ had the lowest average precision (0.63). However, Steak had the lowest average sensitivity (0.13) while ERVcaller had the highest (0.77).

### Analysis of matching short and long-read sequencing samples

We used each tool on six SR-WGS samples and used the matching long-read sequencing data for validation (Table 4). Consistently with the other tests, the tools’ performance varied. Retroseq+ gave the smallest number of predictions; however, 78% of predicted loci were positive for an LTR5_Hs of length >850bp in the corresponding long-read sample. We are particularly interested in the larger insertions as they would suggest a complete LTR (968 bases) that will most likely contain regions of biological importance such as the LTR promoters and enhancers. The great majority of the loci predicted by the tools were confirmed to contain ERV sequences in the long-read data. However, considering only the predicted insertions that correctly contained LTR5_Hs (the target HERV-K element), the performance of the tools varied greatly. For example, 78% of insertions called by Retroseq+ were LTR5_Hs, while only 13% of the Melt calls were LTR5_Hs. All Retroseq+ calls were >850 bases while a substantial proportion of the loci identified by the other tools were smaller. Moreover, the number of predicted insertions also varied, ranging between 18 (Retroseq+) and 481 (Ervcaller). Notably, Steak identified a large number of long LTR5_Hs insertions but over two thirds were reference loci.

**Table 3).**
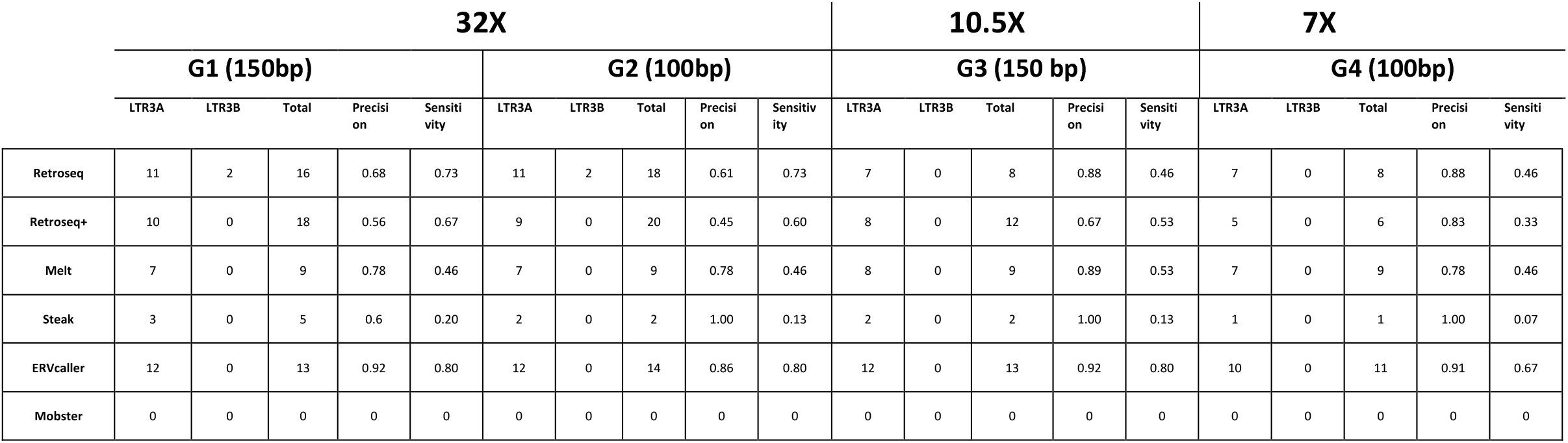
Results of analysis of simulated WGS data. Each row corresponds to a different tool. The table reports the number of correctly identified LTR3A insertions (the target insertions), the number of LTR3B insertions mistakenly classified as LTR3A insertions, and the total number of predicted insertions (Total column) for each sample.

**Table 4).**
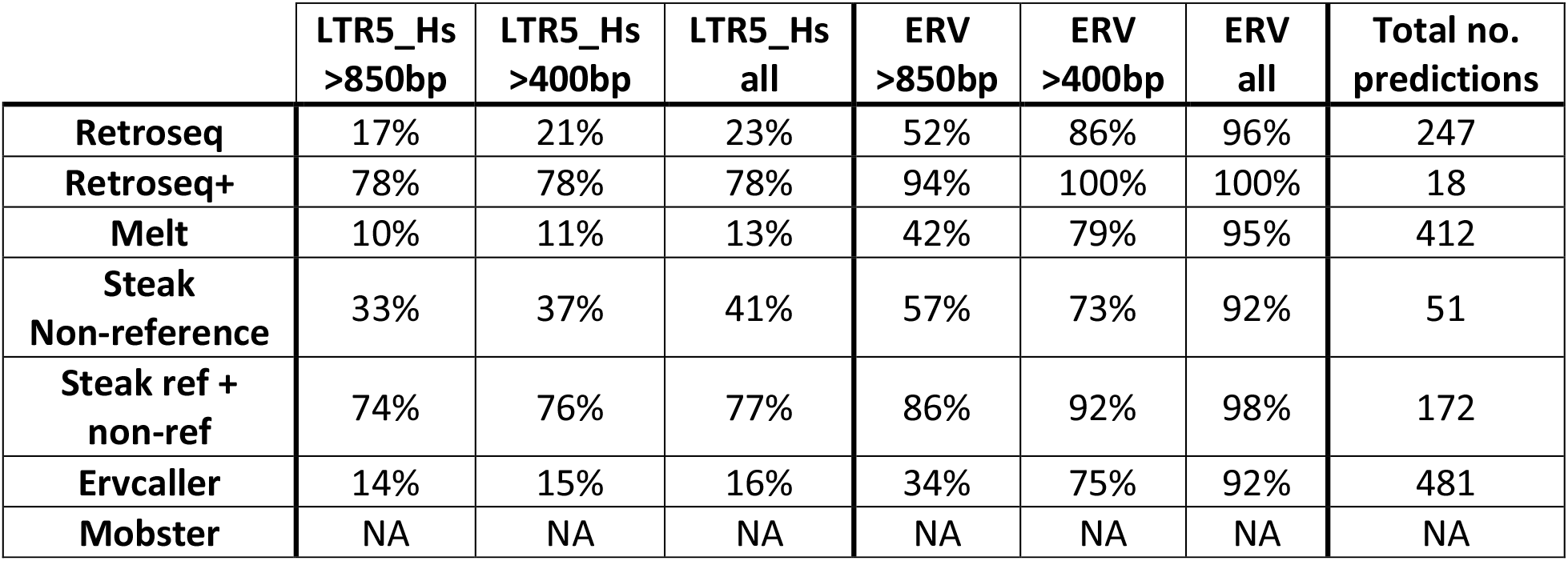
This table shows the proportion of results predicted by each tool in the short-read samples, that are positive for a HERV sequence in the associated long-read data. Each column shows the proportion of results that were HERV-K (LTR5_Hs, the targeted HERV subgroup) or a general HERV sequence and the length of the sequence in the long-read data. The Steak results are reported both including and discarding reference HERV loci. Reference HERV loci were removed for all other tools.

### CPU usage and time

Table 5 shows the CPU Time, Max virtual memory and input file type/size for each tool. Melt had a very low CPU time of 3 hours and 23 minutes; Ervcaller had the highest CPU time of 14 hours and 17 minutes. Mobster usage was not reported as it did not produce any HERV results. Though we used a SAM file with Steak, it is possible to pipe a BAM input into Steak directly. All tools produced intermediate file sizes of a negligible size (<1GB −3 GB) except for Ervcaller which produced intermediate files whose sizes summed to 87 GB. If memory limitations are a concern, tools with minimal CPU time, input file sizes and VM requirements are desirable.

**Table 5).** This table shows how memory is used by each tool. The CPU time is equal to the number of CPUs * time. MAX VM size is the maximum virtual memory used at any one time by any part of the job. The input file size column reports the size of input sequencing data in the format required by each tool. The Max Temporary Files Size shows the maximum temporary storage required by each tool while running. For tools where there is an option to remove (clean up) temporary files, this option was not used.

|  | **CPU Time** | **Max VM Size** | **Input File Format/Size** | **Max Temporary Files Size** |
| --- | --- | --- | --- | --- |
| **Retroseq** | 03:43:01 | 1.15G | BAM / 77G | 2GB |
| **Retroseq+** | 04:01:15 | 1.15G | BAM / 77G | 2GB |
| **Melt** | 03:23:05 | 5.10G | BAM / 77G | 3GB |
| **Steak** | 04:48:35 | 1.17G | SAM / 287G | <1GB |
| **Ervcaller** | 14:17:22 | 22.39G | BAM / 77G | 87GB |

## Discussion and conclusions

This study compared the performance of six computational tools for detecting HERV loci in whole genome sequencing data. Overall, our results provided evidence of their highly variable performance across SR-NGS datasets.

The first test involved applying each tool to 50 WGS samples of real individuals from an ALS cohort. This allowed us to quantify the agreement between tools and the proportion of results that matched to known HERV-K loci. In this experiment Mobster could not identify any HERV insertions confirming its inability to detect this type of elements as was suggested by the authors in their original benchmarking analysis. The agreement between tools ranged between 3% and 63%, and the number of insertions predicted ranged between 296 (Retroseq+) and 13,770 (Steak). A part of this variability can be explained by the fact that Steak was designed to detect the presence of both reference and non-reference HERV insertions, however, the tools’ accuracy might also contribute substantially. Indeed, although 65% of Steak’s predictions matched reference HERV loci, only 1.7% overlapped with the highly characterised, non-reference, polymorphic loci. Looking at the proportion of insertions that matched the non-reference HERV-Ks previously reported in the literature, can inform us about the quality of the predictions made by the tools, as we would expect these to be more likely to be true insertions with respect the novel ones. 39% of Retroseq’s predictions and 52% of Melt’s predictions were previously reported. Ervcaller and Retroseq+ generated sets of predicted loci that greatly matched to previously reported ones (81% and 97.4% respectively, Table 2).

The tools were then applied to simulated WGS. In order to simulate potentially realistic (proviral) insertions, we first generated WGS samples using hg19 varying read length and coverage depth. Then we used a copy of hg19 in which we removed a set of known reference HERV loci, for read mapping of the simulated samples and HERV detection. Therefore, this experiment allowed us to assess how read length and coverage depth affect the tools’ performance and to quantify the tools’ precision and sensitivity. As expected, the tools performed better on high quality WGS data (32X and 150bp reads). Precision and sensitivity highly varied across tools, ranging between 0.56 - 0.92, and 0.20 - 0.80 respectively on the higher quality samples (32X and 150bp reads).

Finally, the tools were tested on six publicly available genomes that had undergone both long and short read sequencing. Given the length of the long reads (>10Kbs), this dataset allowed us to confirm the insertions called in the short-read data using the long reads. In this experiment the great majority (>92%) of all insertions predicted by the tools were confirmed HERVs in the long-read data. However, only Retroseq+ insertions were largely (78%) confirmed to be LTR5_Hs (the target HERV-K element), while the other tools showed a lower ability to distinguish between different HERV LTRs (13-41%). Notably, Steak showed a substantially higher precision for reference loci (>77%) than for non-reference loci (41%)

In conclusion, our analyses showed that tools and protocols developed specifically for the detection of HERV-Ks, such as Ervcaller, Retroseq+, and Steak, generally outperformed generalist tools such as Mobster and MELT. Moreover, the experiments highlighted important characteristics of the tools that the users should consider when designing their analysis pipeline: our implementation of the protocol developed by Wildschutte and colleagues (Retroseq+) produced the most reliable predictions but also the smallest number (296 predictions across 50 genomes); Steak was the only tool able to comprehensively capture the presence of reference HERVs but its performance was substantially higher on reference HERVs than on non-reference HERVs; Ervcaller and Retroseq showed a good balance between number of detected insertions and their quality, however, their performance greatly varied across experiments. For example, they showed high precision and sensitivity in the simulated data, but when applied to real data that is expected to include a large number of other types of insertions (the initial large SR-WGS dataset and the matching short and long read data), both of them showed high sensitivity but low specificity.

Given that all tools presented strengths and weaknesses, we recommend the users to base their choice on the requirements and objectives of the study and to consider combining multiple tools and a consensus approach if computationally feasible. For example, for rare genetic diseases in which both common polymorphisms and rare disruptive variants contribute to their genetics, such as ALS and other neurodegenerative disorders, one could combine the ability of Steak to call reference HERVs, with one of the other tools that showed a higher performance on non-reference insertions. Moreover, according to the availability of biological samples for wet-lab validation, one might choose a more conservative caller such as Retroseq+ or a more sensitive tool such as Ervcaller.

Some of the variability in calling HERV-K insertion sites in our data may stem from the use of the hg19 reference genome. We expect results to differ if hg38 had been used, as it includes more alternate sequences as well as corrections to sequencing artefacts (37). However, the overarching challenge in calling HERVs remains, regardless of reference, as short read sequencing presents intrinsic limitations to capture large insertions. This challenge applies to most types of variants larger than some tens of base pairs and a consensus approach has shown potential, e.g. Gnomad SV (38).

Although long read sequencing can provide a better solution to the detection of large insertions and its use is on the rise, analysing short read sequencing data for large variants is still highly relevant given the great availability of this type of data and its higher sequencing resolution that can allow for a more accurate characterization of variant sequences.

## Supplementary

All of the supplementary data and scripts used for this paper can be found at the following GitHub repository: tools_assessment_hervk_SR-WGS.

## Acknowledgements

This research was supported by the National Institute for Health Research (NIHR) Biomedical Research Centre based at Guy’s and St Thomas’ NHS Foundation Trust and King’s College London. The views expressed are those of the authors and not necessarily those of the NHS, the NIHR or the Department of Health.

## Author Contributions

Conceptualization, AI, CMS, JPQ, AJ and AAC; methodology, AI, HB, RK; software, AI, HB, RK and RD; validation, AI, HB and RK; formal analysis, AI, HB and RK; investigation, AI, HB, RK and CMS; resources, AI and RD; data curation, AI, AAC, AAK, AJ, and RD; writing—original draft preparation, AI, HB, RK; writing—review and editing, CMS, AI, JPQ; visualization, HB and RK; supervision, AI, AAC, CMS, JPQ; project administration, AAC, AI; funding acquisition, AI, AAC. All authors have read and agreed to the published version of the manuscript.

